# Mind the gap: micro-expansion joints drastically decrease the bending of FIB-milled cryo-lamellae

**DOI:** 10.1101/656447

**Authors:** Georg Wolff, Ronald W. A. L. Limpens, Shawn Zheng, Eric J. Snijder, David A. Agard, Abraham J. Koster, Montserrat Bárcena

**Author notes:** Correspondence to Montserrat Bárcena.

## Abstract

Cryo-focussed ion beam (FIB)-milling of biological samples can be used to generate thin electron-transparent slices from cells grown or deposited on EM grids. These so called cryo-lamellae allow high-resolution structural studies of the natural cellular environment by *in situ* cryo-electron tomography. However, the cryo-lamella workflow is a low-throughput technique and can easily be obstructed by technical issues like the bending of the lamellae during the final cryo-FIB-milling steps. The severity of lamella bending seems to correlate with shrinkage of the EM grid support film at cryogenic temperatures, which could generate tensions that may be transferred onto the thin lamella, leading to its bending and breakage. To protect the lamellae from these forces, we milled “micro-expansion joints” alongside the lamellae, creating gaps in the support that can act as physical buffers to safely absorb material motion. We demonstrate that the presence of such micro-expansion joints drastically decreases lamella bending. Furthermore, we show that this adaptation does not create instabilities that could constrain subsequent parts of the cryo-lamella workflow, as we obtained high-quality Volta phase plate tomograms revealing macromolecules in their natural structural context. The minimal additional effort required to implement micro-expansion joints in the cryo-FIB-milling workflow makes them an easy solution against cryo-lamella bending in any biological sample milled on EM grids.

## Main Text

The emergence of focussed ion beam (FIB)-milling has opened an exciting and rapidly growing field in cryo-electron microscopy (cryo-EM) that facilitates structural studies in the natural cellular environment. While small biological specimens (e.g. purified macromolecules) can nowadays be studied at near-atomic resolution by cryo-EM (Cheng, 2018), the limited penetration depth of the electron beam in a transmission electron microscope (TEM) hampers the direct observation of thicker samples like eukaryotic cells. To overcome this limitation, a dual-beam scanning-electron-microscope (SEM) equipped with a FIB and a cryo-sample stage, termed cryo-FIB/SEM, can be used to mill an electron-transparent slice from a cell (Marko et al., 2007; Rigort et al., 2012). Essentially, any internal region of a cell can be accessed by removing excess material above and below it with a focussed beam of gallium ions, resulting in a less than 300 nm thick cryo-lamella that is suitable for *in situ* structural studies of macromolecular complexes by cryo-electron tomography (cryo-ET) (Böck et al., 2017; Bykov et al., 2017; Chaikeeratisak et al., 2017; Engel et al., 2015; Guo et al., 2018; Mahamid et al., 2016).

The FIB-milling workflow to generate lamellae, described in detail in (Medeiros et al., 2018; Schaffer et al., 2015), is very similar across different laboratories and studies, with only small variations tailored to the requirements of specific samples. In short, cells grown or deposited on an EM grid are plunge-frozen and transferred to a cryo-stage in a FIB/SEM instrument. Here, a FIB is applied to the sample at shallow angles, using decreasing beam currents as the distance between the milling areas above and below a region of interest is gradually reduced, and the process in monitored by SEM. The initial higher FIB currents allow the fast removal of bulk material, while low beam currents subsequently ensure a more gentle polishing of the lamella to its final thickness. A layer of organometallic platinum is applied on the sample before milling to further protect the lamellae from unwanted erosion by scattering ions and curtaining (Hayles et al., 2007). Most cryo-lamellae have a width of 10-20 µm and are reproducibly milled to a thickness of 150-300 nm, although special strategies are available as well to produce lamellae below 100 nm thickness, desirable for high resolution reconstructions (Schaffer et al., 2017).

Currently, cryo-FIB-milling is still a rather laborious and time-consuming technique: a day of milling will produce about 5-10 lamellae. These lamellae represent only a tiny fraction of the cellular volume and thus, depending on the specific target to be imaged, may or may not contain the feature of interest. Consequently cryo-FIB-milling is a low-throughput technique and every factor hampering the workflow poses a serious obstacle for its successful application. One of these complicating factors that is regularly observed is the bending of cryo-lamellae during the final milling steps (fig. 1A), which can result in partially or fully broken lamellae (fig. 1B) and usually renders them unusable for subsequent cryo-ET studies.

**Figure 1:**
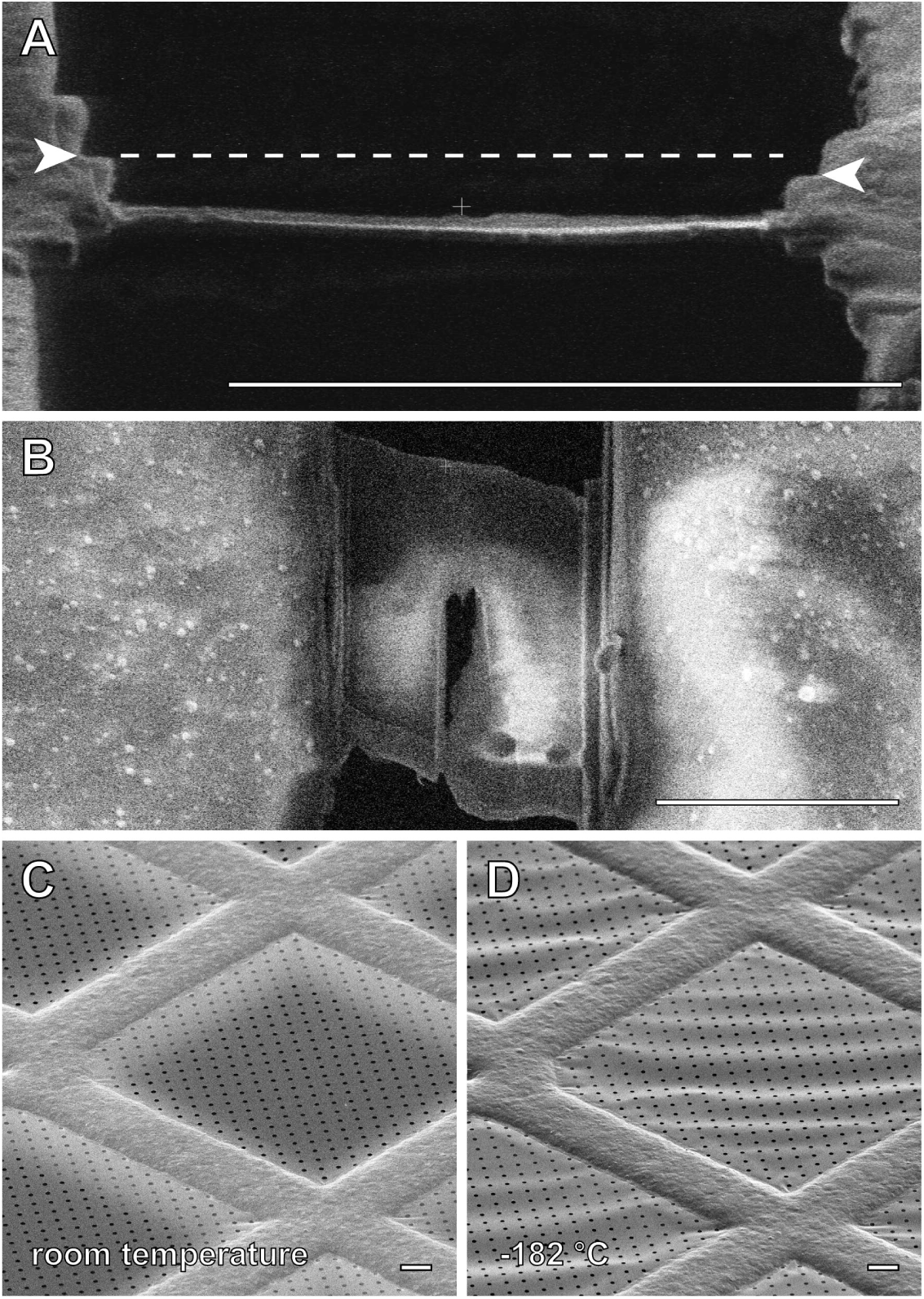
Cryo-lamella bending and temperature stability of the support grids. (A) Bending of a cryo-lamella during the final thinning steps of FIB-milling as observed by FIB-imaging from the direction of milling. The offset between the left and right edge of the lamella (white arrowheads) indicates major sample deformations during milling. (B) Lamella bending leads to unwanted erosion and finally the breaking of the lamella, as observed by SEM-imaging from top view (B). Cryo-lamella bending seems to correlate with the shrinkage of the grid support that occurs during freezing. (C) The grid support film, flat at room-temperature can change to (D) “wavy” at cryogenic temperatures. Scale bars, 10 µm.

While bending usually accounts for a moderate fraction of the total lamellae, in practise it can become a hurdle that is almost impossible to overcome. In particular, we have noticed that the FIB-milling success rate can critically depend on the specific batch of commercial grids commonly used in the field, whose quality is not directly under the control of the researcher. Ideally, the desired grid support should be flat at all temperatures; however, it has been observed that its morphology can change at low temperatures due to shrinkage (fig. 1C-D), resulting in a “wavy” support film. This waviness correlates with an increase in lamella bending, likely because the deformations of the support that arise during milling generate tensions in the sample (M. Schaffer, personal communication). We hypothesized that, if these tensions are released from the grid onto the cryo-lamellae during milling, generating “micro-expansion joints” alongside a lamella could protect it from such forces and thus prevent their bending. The concept of expansion joints originates from construction engineering: structures, such as concrete floors, are equipped with gaps to provide a physical buffer that will safely absorb any developing motion. In our case, they are intended to absorb relaxations of the grid support released upon milling, which otherwise would harm the integrity of the cryo-lamella.

We developed an improved protocol, which includes the milling of lateral micro-expansion joints into the grid support flanking each lamella site (fig. 2A-D) and reduces lamella bending. The additional milling time is minimal (2-3 min per lamella) as the micro-expansion joints are milled with the initial high currents and are only ∼1 µm wide. They cover the same length as the area milled for lamella generation (in our case about 50 µm) and are at a distance of 3-4 µm from the lamella, leaving a sufficient amount of cellular material to support the lamella. Once micro-expansion joints are applied, relaxations in the grid support next to them often become apparent (see suppl. fig. 1), pointing towards a release of physical tensions which otherwise probably would have caused lamella bending during thinning. Furthermore, just a single micro-expansion joint can already improve lamella stability in case a milling site is too close to a grid bar. Subsequent milling steps remain unchanged and thus this adaptation can easily be incorporated in any cryo-lamella protocol. In our case, milling of 10 µm-wide lamellae was performed in four steps using FIB currents from 1 nA to 30 pA (see supplemental data for details), with a final polishing step thinning them to 130–230 nm thickness. Additionally, we used an increased milling angle during initial rough milling (16° relative to the grid surface) and gradually reduce it to 11° during polishing. Using a higher angle at the initial milling step allows for better visualization and interpretation of the sample topology. Moreover, this seemed to help avoid excessive abrasion of the organometallic platinum on the front end of a cryo-lamella, which in our experience increases with lower milling angles, especially when using higher beam currents.

**Figure 2:**
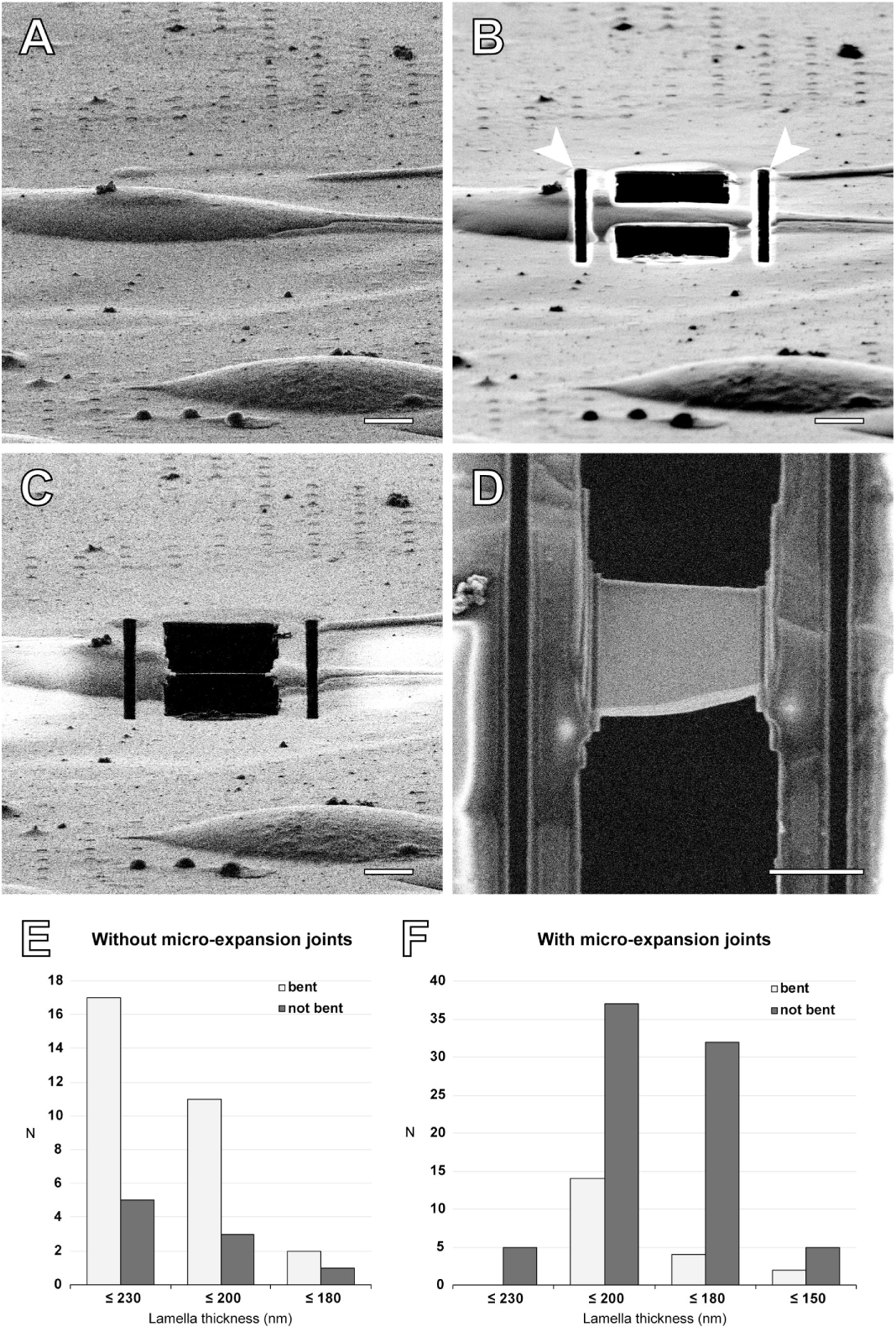
Incorporation of micro-expansion joints into the cryo-FIB-milling workflow. (A) First, eukaryotic cells (murine 17clone1 cells in these images) are grown and plunge-frozen on EM grids, and suitable cells for cryo-FIB-milling are selected. (B) In addition to the two rectangular milling patterns above and below the region of interest, lateral micro-expansion joints (white arrowheads) are milled with higher ion-beam currents (1 – 0.5 nA) during the initial rough milling step. They are at a distance of 3-4 µm from the lamella, leaving a sufficient amount of cellular material to support the lamella. (C) Subsequently, the lamella is gradually thinned while reducing the ion-beam currents down to 30 pA during polishing to the final thickness. (A-C) From the ion-beam view the sample morphology and the lamella thickness can be assessed during the process. (D) SEM top-view image of the final lamella showing the typical homogeneous contrast of a thinned and fully intact lamella without signs of bending. (E,F) Comparison of the success rates in the milling of all the cryo-lamellae prepared in this study. (E) When micro-expansion joints were not applied to the lamella site, the majority of the lamellae bent during the final milling steps, regardless of the intended thickness (77% bent lamellae in total). (F) Upon addition of micro-expansion joints, almost 80% of the lamellae could successfully be thinned to their final thickness without signs of bending, which allowed the reproducible generation of lamellae down to a thickness of 130 nm (see also fig. 3).

In order to assess the impact of this adaptation on the overall success rate of the cryo-lamella workflow, we generated a series of lamellae either with or without micro-expansion joints using EM grids of a batch showing the “wavy” support film properties described above. To rule out any influence of variations in quality per grid, both cryo-lamellae with (43 in total) and without (27 in total) micro-expansion joints were milled on each grid (9 grids in total), resulting in a total of 70 lamellae with a thickness of 170-230 nm. While for lamellae without micro-expansion joints only 22% could be thinned to their final thickness without bending, we were able to successfully mill 77% of the lamellae when applying micro-expansion joints. This demonstrated that the addition of micro-expansion joints results in a highly significant increase in the success rate of cryo-lamella milling (*p*-value 1.2 × 10^-5^, chi-square test), which encouraged us to further reduce the thickness of the lamellae generated by this protocol. We were able to reliably mill cryo-lamellae down to a thickness of 130 nm, which extended our data to a total of 139 lamellae (see fig. 2 E-F, fig. 3).

**Figure 3:**
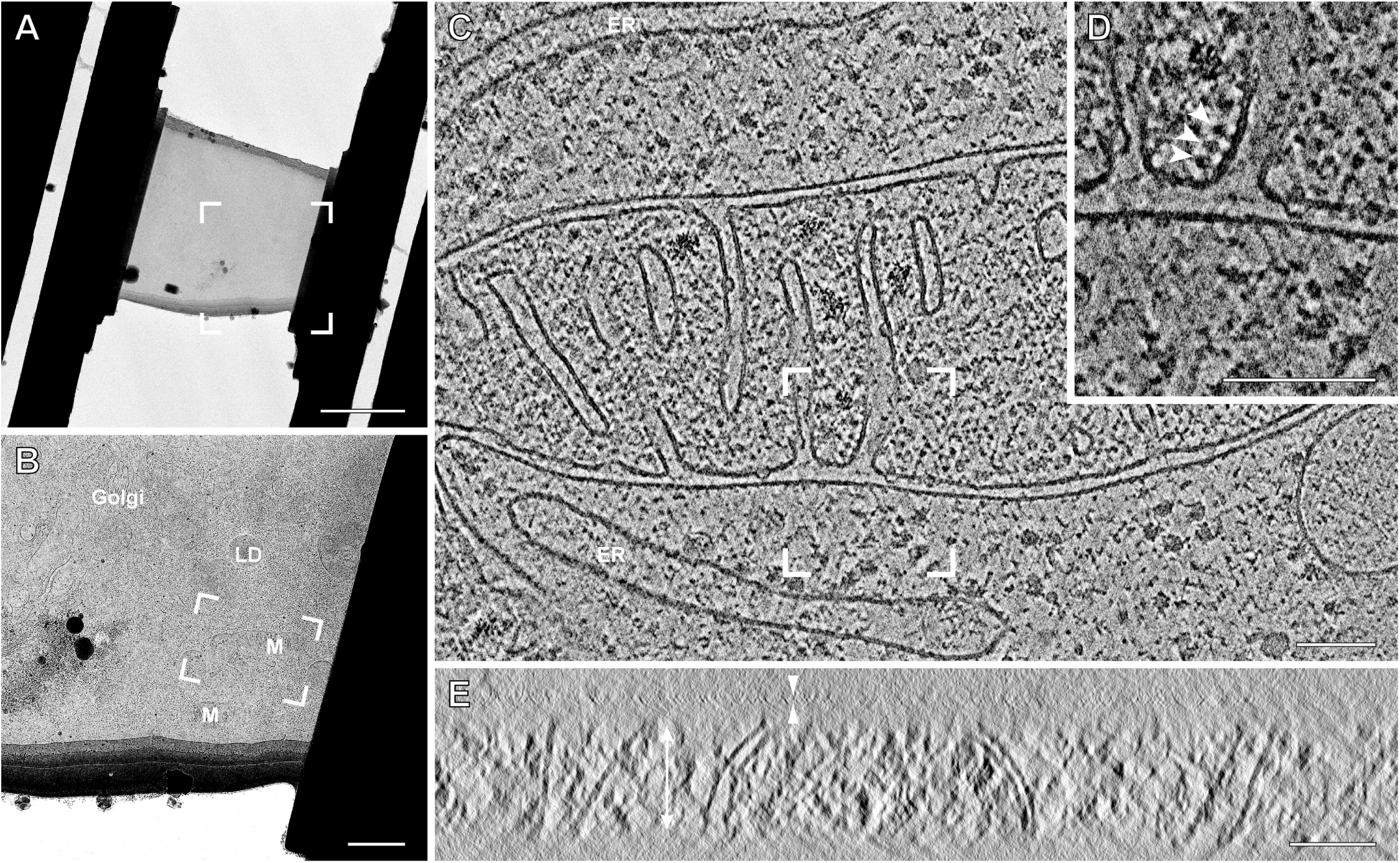
TEM imaging and cryo-electron tomography on a lamella milled with micro-expansion joints. (A) A cryo-lamella with micro-expansion joints observed in a 300 kV TEM. The low-resolution overview shows the structural integrity of the lamella. The boxed area indicates the region enlarged for identification of intracellular membranous structures (B), such as the Golgi apparatus (Golgi), lipid droplets (LD) or mitochondria (M). From the selected area of interest (white box), a tilt-series was acquired using a Volta phase plate. (C) Slice through a tomogram of this area, reconstructed with AreTomo (see supplementary material), which shows a mitochondrion, endoplasmic reticulum (ER) membranes and various macromolecules in their structural context. (D) A close-up of the mitochondria allows the identification of individual ATP-synthase complexes embedded in the cristae membranes (white arrowheads). (E) A x,z slice of the tomogram makes apparent (from top to bottom) the ∼5 nm thick Pt layer on top of the lamella (white arrowheads), a ∼20 nm thick intermediate layer of condensed water accumulated between the final milling step and the platinum sputtering, and the 130 nm thick lamella (double headed arrow). Tomographic data collection was done at -0.5 µm defocus and a pixel size of 0.351 nm. Scale bars, 5 µm (A), 1 µm (B) and 100 nm (C-E).

While the generation of micro-expansion joints clearly prevented lamella bending, the removal of additional material from the cell that holds the hanging lamella and from the underlying support could potentially create instability and conductivity issues affecting other parts of the workflow, namely, transfer and imaging. To explore these concerns we imaged a major fraction of the cryo-lamellae with micro-expansion joints in a 300 kV TEM (fig. 3). The transfer of lamellae between instruments is one of the most critical steps in the cryo-lamella workflow, which can easily lead to the destruction of the lamellae. Of the 56 lamellae imaged, 55 did not show any transfer damage while only one had a crack. Therefore, the addition of micro-expansion joints does not seem to reduce the stability of the lamellae during transfer. To assess whether micro-expansion joints would be limiting the stability and conductivity during TEM imaging, we performed cryo-ET using the Volta phase plate (an imaging mode particularly sensitive to charging effects (Danev and Baumeister, 2017; Mahamid et al., 2016)) on suitable areas containing recognizable cellular features. Our experimental data indicates that micro-expansion joints, even though they interrupt part of the conductive layer between the lamella and the EM grid, do not hamper Volta phase plate imaging as we were able to collect automated tilt-series without signs of charging. Furthermore, we reconstructed high quality tomograms containing mitochondria (fig. 3C-E) in which individual macromolecular structures could be identified (e.g. ATP-synthase complexes embedded in the cristae membranes of mitochondria (Davies et al., 2014), fig. 3D).

In summary, our results demonstrate a highly significant increase in the success rate of cryo-lamella preparation by adding micro-expansion joints to milling sites on difficult support films without affecting their stability during transfer and imaging. The effectiveness of this approach strongly supports the notion that deformations in the grid support are a major cause for lamella bending, a reoccurring problem in cryo-FIB-milling that currently limits the yield of this technique. We showed that it is possible to perform cryo-ET using the Volta phase plate on these lamellae, which poses one of the most demanding ways of cryo-ET (Danev and Baumeister, 2017). With the minimal effort required to include micro-expansion joints in any cryo-lamella protocol, this method can easily increase the throughput for every sample type milled on EM-grids.

## Acknowledgements

We thank Miroslava Schaffer (MPI Martinsried, Germany) for her helpful discussions and advice. High resolution EM data was collected at The Netherlands Centre for Electron Nanoscopy (NeCEN) with assistance from Christoph Diebolder.

