## Supplementary Material for "Mind the gap: micro-expansion joints drastically decrease the bending of FIB-milled cryo-lamellae"


#### Author names and affiliations:

Georg Wolff[1], Ronald W. A. L. Limpens[1], Shawn Zheng[2], Eric J. Snijder[3], David A. Agard[2], Abraham J. Koster[1] & Montserrat Bárcena[1*]

[1] Department of Cell and Chemical Biology, Section Electron Microscopy, Leiden University Medical Center, Leiden 2333 ZC, The Netherlands

[2] Department of Biochemistry & Biophysics, University of California, San Francisco, California, USA

[3] Department of Medical Microbiology, Molecular Virology Laboratory, Leiden University Medical Center, Leiden 2333 ZA, The Netherlands

### Supplementary material

#### Materials and methods

**Cell culture and vitrification.** R1/4 gold Quantifoil grids, (200 mesh, Quantifoil Micro Tools, Jena, Germany) were cleaned overnight by placing them on a chloroform soaked filter paper in a sealed glass dish. After letting them dry, the grids were glow-discharged and placed in 3.5 cm diameter cell culture dishes (Greiner Bio-One, Frickenhausen, Germany) containing PBS. After UV-sterilization for 30 min in a laminar-flow, PBS was replaced by cell culture medium containing 100,000 17clone1 or VeroE6 cells per dish and incubated at 37 °C overnight. After retrieval of the grids from the cell culture dish using 5/15 style kinked tweezers (Dumont, Montignes, Switzerland), they were transferred to the tweezers of a Leica EM GP (Leica microsystems, Wetzlar, Germany) automated plunge-freezer, and plunge frozen after applying 15 s of blotting from the backside of the grid with chamber conditions of 37 °C and 95% humidity. After being plunge frozen in liquid ethane, the grids were stored in liquid nitrogen until used for cryo-FIB-milling.

**Cryo-FIB-Milling.** Lamella preparation was done in an Aquilos cryo-FIB/SEM (Thermo Fisher Scientific, Hillsboro, OR, USA). First, each grid was sputter-coated with platinum (1 kV, 30 mA, 10 Pa, applied for 10 s) to increase conductivity. Subsequently a layer of organometallic platinum was applied onto the sample by the gas injection system (GIS) operated at 28 °C for 6 - 7 seconds. Four milling steps (rough, intermediate, fine, polishing) were performed per lamella, for which a 30 kV ion beam was used with 1 to 0.5 nA, 300 to 100 pA, 50 pA and 30 pA probes, respectively. The distance between the milling patterns was 3-5 µm during rough milling, 1-1.2 µm during the intermediate step, and 400-500 nm during fine milling. In the rough and intermediate milling the angle was set to 16° and 12°, respectively. During polishing the cryo-lamellae were thinned to 130–230 nm at a FIB angle of 11° relative to the grid surface. After polishing all the lamellae per grid, the sample was sputter-coated with a ~5 nm thick layer platinum (1 kV, 10 mA, 10 Pa, applied for 4 s) with the integrated sputter coater. The process was monitored by FIB and SEM imaging. SEM imaging was done at 3 kV with either a 13 or 25 pA probe with dwell times of 100–500 ns, while FIB imaging was performed at 30 kV using a 10 pA probe with a dwell time of 100 ns.

**Cryo-ET data collection.** Tilt-series acquisition was carried out on a FEI Titan Krios (Thermo Fisher Scientific, Hillsboro, OR, USA) operated at 300 kV and equipped with a Gatan GIF Quantum energy filter (Gatan, Pleasanton, CA). Movie-frames were collected using a Gatan K2 Summit direct detection camera operated in counting mode. The imaging conditions were as follows: EFTEM microprobe, 42,000x magnification, pixel size of 3.51 Å, zero-loss imaging with a 20 eV slit width, beam diameter 3 µm, defocus -0.5 µm. Bi-directional tilt-series acquisition was performed with FEI Tomo4 software, started at 0° with 2° increments and ranged from -50° to +70° stage tilt, to account for the fact that the lamella surfaces were orthogonal to the electron beam at 11°. Volta phase plate was activated before each tilt-series acquisition until the phase shift stabilized at approximately 0.5 pi. The total dose applied on the sample was 140 $e^-$ per $Å^2$ spread over 61 tilts with a dose distribution factor of 1.6, resulting in an average exposure time of 2.8 s per tilt.

**Alignment and reconstruction.** After data collection, the movie frames per tilt (14 each) were aligned with MotionCor2 (Zheng et al., 2017). Tilt-series alignment and tomogram reconstruction were performed with AreTomo. Developed at University of California San Francisco (Zheng and Agard, to appear) and implemented on a multi-GPU Linux platform, this program fully automates the tomographic workflow from fiducial-free alignment to SART (Simultaneous Algebraic Reconstruction Technique) reconstruction. A typical tomogram can be obtained within a few minutes on a platform with two NVIDIA GeForce GTX 780 Ti GPUs.

#### Supplementary figures

**Figure S1:** Motions in the grid support become apparent upon micro-expansion joint milling. (A) A eukaryotic cell (murine 17clone1 cell) grown and plunge-frozen on an EM grid is selected for cryo-lamella FIB-milling. (B) Upon milling, motion in the support film next to one of the micro-expansion joints becomes apparent (white arrowhead). These deformations are probably caused by tensions in the grid support, that otherwise could have harmed the integrity of the lamella.
