## Supplementary figures and images for "Mind the gap: micro-expansion joints drastically decrease the bending of FIB-milled cryo-lamellae"

### Supplementary Figure 1

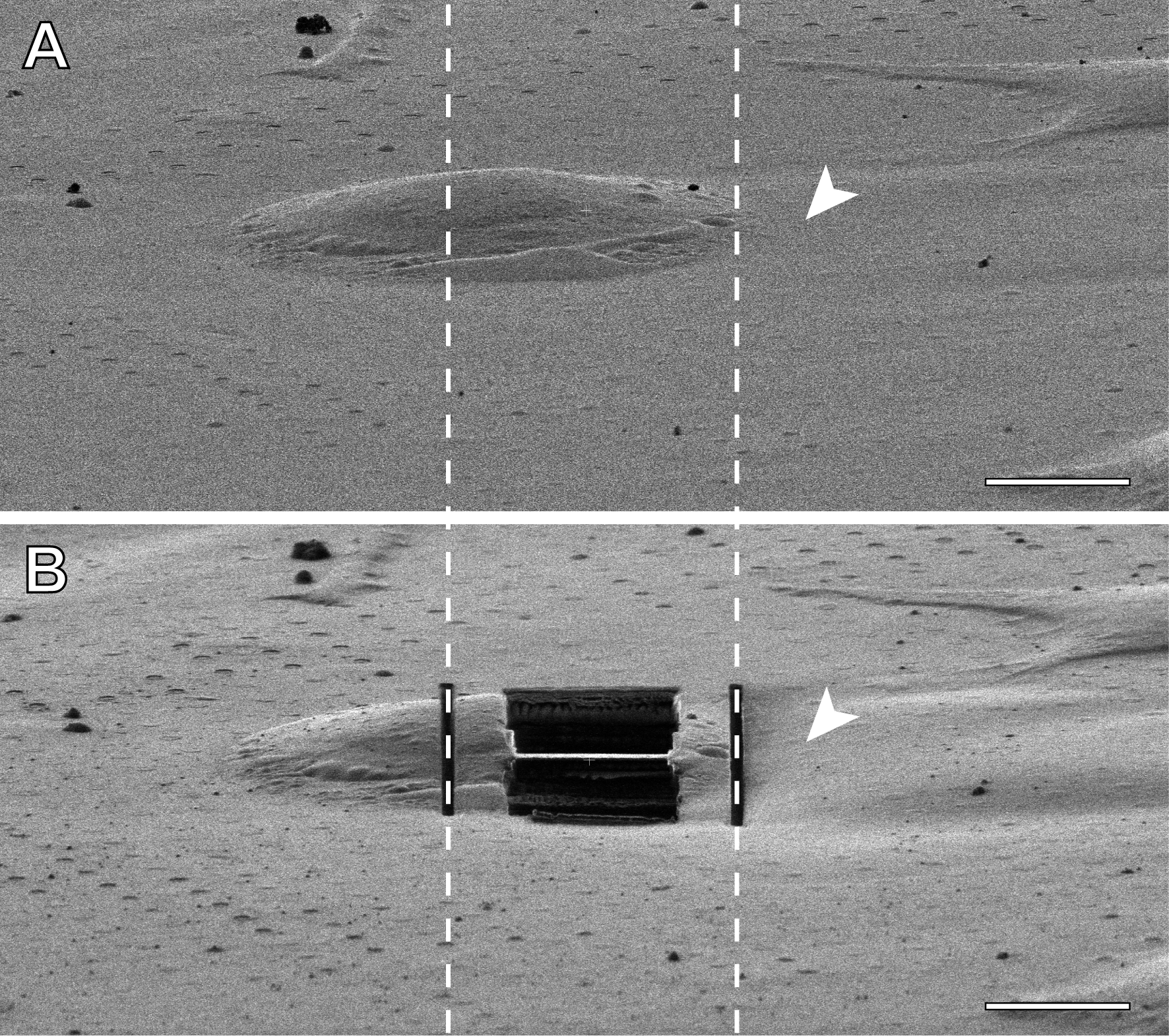
